# Deciphering the interstrand crosslink DNA repair network expressed by *Trypanosoma brucei*

**DOI:** 10.1101/543975

**Authors:** Ambika Dattani, Shane Wilkinson

**Affiliations:** School of Biological and Chemical Sciences, Queen Mary University of London, London, E1 4NS UK

**Author notes:** Current address: School of Life Sciences, Queen’s Medical Centre, University of Nottingham, Nottingham, UK.

## Abstract

Interstrand crosslinks (ICLs) represent a highly toxic form of DNA damage that can block essential biological processes including DNA replication and transcription. To combat their deleterious effects all eukaryotes have developed cell cycle-dependent repair strategies that coopt various factors from ‘classical’ DNA repair pathways to resolve such lesions. Here, we report that *Trypanosoma brucei*, the causative agent of African trypanosomiasis, possesses such systems that show some intriguing differences to those mechanisms expressed in other organisms. Following the identification of trypanosomal homologues encoding for CSB, EXO1, SNM1, MRE11, RAD51 and BRCA2, gene deletion coupled with phenotypic studies demonstrated that all the above factors contribute to this pathogen’s ICL REPAIRtoire with their activities split across two epistatic groups. We show that one network, which encompasses TbCSB, TbEXO1 and TbSNM1, may operate throughout the cell cycle to repair ICLs encountered by transcriptional detection mechanisms while the other relies on homologous recombination enzymes that together may resolve lesions responsible for the stalling of DNA replication forks. By unravelling and comparing the *T. brucei* ICL REPAIRtoire to those systems found in its host, targets amenable to inhibitor design may be identified and could be used alongside trypanocidal ICL-inducing agents to exacerbate their effects.

**Author summary:** Parasites belonging to the *Trypanosoma brucei* complex cause a human and animal infections collectively known as African trypanosomiasis. Drugs used against these diseases are problematic as medical supervision is required for administration, they are costly, have limited efficacy, may cause unwanted side effects while drug resistance is emerging. Against this backdrop, there is a need for new therapies targeting these neglected tropical diseases. Previous studies have shown compounds that induce DNA interstrand crosslinks (ICLs) formation are effective trypanocidal agents with the most potent invariably functioning as prodrugs. Despite the potential of ICL-inducing compounds to treat African trypanosomiasis little is known about the ICL repair mechanisms expressed by trypanosomes. Using a combination of gene deletion and epistatic analysis we report the first systematic dissection of how ICL repair might operate in *T. brucei*, a diverged eukaryote. It sheds light on the conservation and divergence of ICL repair in one of only a handful of protists that can be studied genetically, and offers the promise of developing or exploiting ICL-causing agents as new anti-parasite therapies. These findings emphasise the novelty and importance of understanding ICL repair in *T. brucei* and, more widely, in non-model eukaryotes.

## Introduction

Spread by the hematophagous feeding behaviour of tsetse flies, protozoan parasites belonging to the *Trypanosoma brucei* species complex are responsible for a group of human and animal infections collectively known as African trypanosomiasis [1]. In terms of their medical importance, these pathogens have caused several major epidemics across sub-Saharan Africa with the last ending at the turn of the millennium [2]. Based on concerted efforts by the World Health Organisation (WHO), national control programmes and non-governmental organisations, improved screening and new treatment approaches have been implemented resulting in a drastic reduction in the estimated number of cases of human African trypanosomiasis (HAT) from around 300,000 in 1998 to below 20,000 in 2015 [1]. Based on the sustained success of such schemes WHO aims to eliminate the human form of the disease as a public health problem by 2020, a goal that is now tantalisingly close [3, 4].

Despite significant progress being made to eradicate HAT, a potential drawback surrounds the therapies being used as each have their own problems; some have limited efficacy, others elicit unwanted side effects, most require prolonged periods of hospitalisation while many are expensive to synthesize, transport and/or store [5, 6]. Additionally, parasite strains refractory to treatment have been generated in the laboratory with some of the resistance mutations noted in these lines also observed in the field [7–12]. Together these issues could potentially derail WHOs ambition for HAT elimination. One approach that may help overcome such problems involves determining the mechanism of action for these existing therapies to exploit any findings in the development of new trypanocides. Applying this strategy to nifurtimox, one of the components in the clinically used nifurtimox eflornithine combination therapy, revealed that a mitochondrial NADH dependent type I nitroreductase, an enzyme commonly found in bacteria and some lower eukaryotes but absent from higher eukaryotes, plays a key role in catalysing the conversion of the 5-nitrofuran prodrug to its cytotoxic products [13]. Screening programmes utilizing this activity has identified a range of anti-T. *brucei* compounds. This revealed that chemicals containing functional groups that can promote DNA damage *via* formation of cross-linkages display significant potency against the bloodstream form of this parasite while exhibiting low toxicity to mammalian cells [14–17].

Generated by endogenous metabolic processes and exogenous mutagens, interstrand crosslinks (ICLs) represent a particularly dangerous type of DNA damage. Formed when the complementary strands within the DNA double helix become covalently linked, they block essential cellular processes that require strand separation such as DNA replication and transcription. If left unchecked, they can go on to promote chromosomal fragmentation, rearrangements, or cell death [18–21]. To conserve genome integrity, all cells exploit various combinations of enzymes from “classical” DNA repair pathways to help resolve ICL damage with these constituting an organism’s so-called ICL REPAIRtoire. The precise mechanism that repair ICLs are unclear as different systems predominate at different stages in the cell cycle while evolutionarily diverse organisms employ distinct mechanisms to accomplish this task [22–24]. In transcriptionally active mammalian cells, the ICL-mediated stalling of an RNA polymerase complex results in recruitment of factors such as CSB to the lesion site with these subsequently interacting with other DNA repair enzymes including SNM1A, XPG and XPF-ERCC1 [25, 26]. In a process known as ‘unhooking’, the XPF-ERCC1 and XPG endonucleases cleave either side of the crosslinked sugar-phosphate backbone in one of the DNA strands with nucleases, including SNM1A, subsequently degrading the released sequence up to and beyond the ICL [27]. The resultant gap is then filled using damage tolerant translesion synthesis (TLS) DNA polymerases such as Pol *ζ*, Pol *η*, Pol *ι*, Pol *Θ* or Pol *κ* with DNA ligase restoring the double strand DNA (dsDNA) structure [28–30]. Once the above has taken place, a second round of nucleotide excision repair (NER), TLS DNA polymerase and DNA ligase activities completely removes the unhooked ICL from the second DNA strand.

In dividing mammalian cells, the ICL-mediated stalling of a single or converging DNA replication fork(s) are recognised by the FANCM helicase. This promotes reversal of the replication fork(s), possibly involving polyubiquitination of PCNA by RAD5-like (e.g. HLTF) activities [31, 32], and recruits a large multi-subunit ubiquitin ligase, termed the Fanconi Anaemia (FA) core complex (FANCA, −B, −C, −E, −F, −G, −L and −T plus the ancillary factors FAAP20, −24 and −100), to the site of the DNA lesion [33–36]. Once formed, this complex monoubiquitinates FANCD2/FANCI with the activated heterodimer drafting a series of nucleases (e.g. FANCQ (XPF)-ERCC1, MUS81-EME1, FAN1 and FANCP (SLX4)) to the site of DNA damage [35, 37]. These cleave the DNA backbone at sequences 5’ and 3’ to the lesion resulting in the unhooking of the ICL from one of the DNA strands, formation of a single stranded gap and generation of dsDNA breaks (DSBs). The single stranded gap is filled and integrity of the sugar-phosphate backbone is restored by TLS DNA polymerase(s) and DNA ligases activities. This links the parental ICL-containing DNA molecule to one of the newly synthesised DNA strands. The ICL is then completely removed by the concerted action of NER, TLS DNA polymerase and DNA ligase activities. The DSB is recognised by the MRN complex (MRE11/RAD50/NBS1) that guide components of the homologous recombination (HR) pathway (*e.g*. FANCD1 (BRCA2), FANCR (RAD51), FANCO (RAD51C)) to mediate recombination between the broken dsDNA molecule and the newly repaired DNA structure generated from the second round of NER/TLS DNA polymerase/DNA ligase activities. This results in reformation of the Y-shaped forked structure that can serve as a template for recommencement of DNA replication [38]. Due to the bidirectional nature of DNA replication, ICLs can cause the stalling of two converging DNA replication forks [39, 40]. Such X-shaped structures can be resolved using the mechanism outlined above although in this case two of the other newly synthesised DNA strands both form DSBs. In this situation, the above HR mediated repair systems are activated with the newly repaired DNA strand serving as template. The outcome of this event is formation of two dsDNA molecules and the disassociation of the replication machinery.

Little is known about the ICL repair mechanisms expressed by trypanosomes despite these parasites being highly susceptible to ICL-inducing compounds. Some components of the global genome NER (GG-NER) pathway (e.g. TbXPC and TbDDB) may potentially be involved in this process [41] while *T. brucei* cells null for TbSNM1, a member of the SNM1/PSO2 family of nucleases, display sensitivity only to ICL-inducing compounds [42]. Here, using a classical genetics-based approach, we analyse whether other DNA repair enzymes from the parasite’s transcription-coupled NER (TC-NER) (TbCSB), HR (TbMRE11, TbRAD51, TbBRCA2 and TbEXO1), mismatch repair (MMR; TbEXO1) and DNA damage tolerance (TbREV3 and TbRAD5) pathways contribute to the trypanosomal ICL repair network then assess their interplay with TbSNM1 and with each other.

## Results

### Identifying putative trypanosomal ICL repair enzymes

Previous studies have shown that MRE11, CSB, EXO1, RAD5 and REV3 all contribute to the yeast and/or mammalian ICL repair systems [25, 26, 43–45]. Using reciprocal BLAST the *T. brucei* genes encoding for homologues of these enzymes plus their flanking regions were identified from the TriTrypDB (http://tritrypdb.org/tritrypdb/) and NCBI (https://www.ncbi.nlm.nih.gov/) databases. For TbMRE11 and TbCSB, the sequences arising from these searches corresponded to those previously reported [41, 46, 47]. In contrast, the genes described here as encoding EXO1, RAD5 and REV3 orthologues have not been characterised. The 2394 bp hypothetical gene (Gene ID: Tb927.4.1480) assigned here as Tbexo7 has potential to encode for a 86.7 kDa enzyme (TbEXO1) that shares approximately 24 % sequence identity with EXO1/HEX1 exonucleases. The *T. brucei* homologue contains within its amino terminal the XPG_N (PF00752) and XPG_I (PF00867) domains, that at this position in the protein sequence is a characteristic of this family of enzyme (Fig 1A). A 4839 bp hypothetical open reading frame (Gene ID: Tb927.7.1090), designated here as *Tbrad5*, was identified as having potential to encode for a homologue of human helicase-like transcription factor (HLTF) and *Saccharomyces cerevisiae* RAD5. The estimated 179 kDa trypanosomal enzyme (TbRAD5) has approximately 24 % sequence identity to this family of DNA helicase/ubiquitin ligases. Homology is centred on domains responsible for helicase (SNF2_N (PF00176); Helicase_C (PF00271)) and ubiquitin ligase (zf-RING_2 (PF13639)) activities, and DNA binding (HIRAN (PF08797)) (Fig 1B) with the order in which these regions are arranged in the protein backbone being conserved between the parasite, yeast and fungal homologues. A 6181 bp hypothetical gene (Gene ID: Tb927.8.3290; designated as Tbrev3) was postulated to encode for a 217 kDa protein (TbREV3) that has approximately 25 % identity to the error-prone DNA polymerase ζ catalytic subunit expressed in *S. cerevisiae* and human cells. The parasite enzyme has a domain structure typical of REV3 subunits containing a DNA polymerase family B, exonuclease domain (PF03104) towards its amino terminal and a DNA polymerase family B (PF00136) followed by C4-type zinc finger (PF14260) towards its carboxyl terminal (Fig 1C).

**Figure 1:**
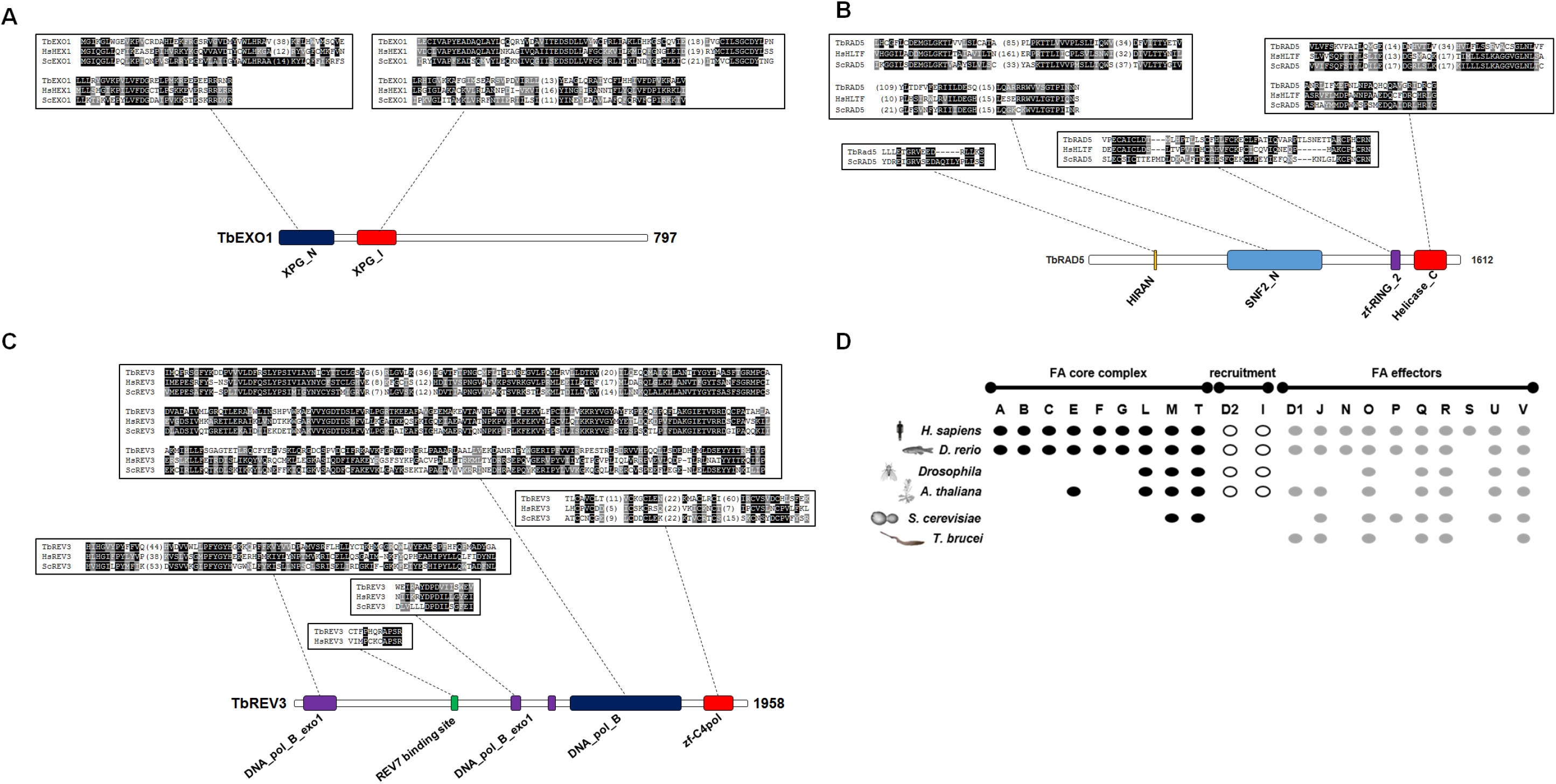
Sequence analysis of potential trypanosomal ICL repair enzymes. **(A)**. Sequences corresponding to the XPG N-terminal (XPG_N) and XPG_I protein (XPG_I) domains of TbEXO1 (accession number XP_847153) were aligned with the equivalent regions from the *H. sapiens* (HsHEX1; AAC32259) and *S. cerevisiae* (ScEXO1; NP_014676) orthologues. **(B)**. Sequences corresponding to the HIRAN, SNF2 family N-terminal (SNF2_N; blue box), RING finger (zf-RING_2; purple box) and helicase conserved C-terminal (Helicase_C; red box) domains of TbRAD5 (XP_845733) were aligned with the equivalent regions from the *H. sapiens* (HsHLTF; NP_001305864) and *S. cerevisiae* (ScRAD5; NP_013132) orthologues. **(C)**. Sequences corresponding to the DNA polymerase family B, exonuclease (DNA_pol_B_exo1), DNA polymerase family B (DNA_pol_B) and C4-type zinc-finger of DNA polymerase delta (zf-C4pol) domains, and to the putative REV7 binding site of TbREV3 (XP_847160) were aligned with the equivalent regions from the *H. sapiens* (HsREV3; NP_002903) and *S. cerevisiae* (ScREV3; CAA97873) orthologues. In the alignments, conserved residues are highlighted in black or grey. **(D)**. Comparison of the Fanconi Anaemia (FA) pathway components expressed by *Danio rerio* (*D. rerio*; fish), *Drosophila melanogaster* (*Drosophila*; insect), *Arabidopsis thaliana (A. thaliana*; plant), *Saccharomyces cerevisiae* (*S. cerevisiae*; yeast) and *T. brucei* in relation to the system possessed by humans *(H. sapiens)*. Trypanosomal FA factors were identified by reciprocal BLAST analysis using human Fanconi anemia (FA) factors as bait. An oval indicates the possibility for the presence of a given factor whereas no oval indicates that that component is most probably absent. In mammalian cells, the FA ICL repair pathway can be subdivided the FA core complex (black ovals), recruitment factors (white ovals), FA effectors (grey ovals). In addition to lacking sequences corresponding to the FA core (FANCA, −B, −C, −E, −F, −G, −L and −T) or recruitment (FANCD2A and −I) complexes, no discernible *T. brucei* homologues for many of the FA associated proteins (FAAP10, −10, −20, −24 and −100) were detected. Figure layout derived from Dong et al., 2015 [49].

In humans, the FA pathway plays a major role in resolving ICLs encountered during DNA replication. It is now recognised that other eukaryotes also express this system although the activities involved vary between species [48, 49]. To determine the extent of this system in *T. brucei*, FA proteins (complete or specific domains) from humans, *Danio rerio, Drosophila melanogaster, Arabidopsis thaliana* and *S. cerevisiae* were used to search for trypanosomal homologues held on the TriTryp and NCBI databases. Out of the 26 sequences examined, *T. brucei* possessed discernible homologues for 7 including BRCA2 (FANCD1), RAD51 (FANCR), RAD51C (FANCO) and XPF (FANCQ), enzymes that play a role in this parasite’s HR and NER pathways (Fig 1D) [41, 50–52]. All of the sequences identified were so-called FA effector proteins with none corresponding to components of the FA core complex (FANCA, −B, −C, −E, −F, −G, −L, −M and −T), recruitment factors (FANCD2A and −I) or FA associate protein (FAAP10, −16, −20, −24 and −100).

### Construction and validation of *T. brucei* single null mutant lines

To evaluate whether the identified sequences were involved in the parasite’s ICL repair network, cells null for each gene were generated. To achieve this, DNA fragments containing the 5’ or 3’ flanking or coding sequences from the gene of interest were amplified from wild type *T. brucei* genomic DNA, digested with restriction enzymes (5’ sequences were processed with SacI and XbaI while 3’ sequences were treated with Apal and Kpnl) then cloned either side of a drug resistance cassette that contains the gene encoding for the hygromycin B phosphotransferase *(hyg)* or neomycin phosphotransferase (neo) (S1A Fig). The gene deletion/disruption constructs were linearized with SacI/KpnI and the fragments transformed into bloodstream form *T. brucei*. To remove both allelic copies of the target gene (all sequences tested were single copy genes) two rounds of nucleofection were performed to firstly create heterozygous and then null mutant lines, with clonal populations of the latter tentatively designated as *T. brucei* Tb*csb*Δ, Tb*mre11*Δ, Tb*exol*Δ, Tb*rad5*Δ and Tb*rev3*Δ. The predicted effect of each gene disruption event with each integration fragment is depicted (S1B-G Figs).

To confirm that integration of the introduced DNA fragments had occurred into the correct genetic loci, DNA amplification reactions using gDNA as template were performed. Such PCRs used primer combinations that generated fragments specific for the intact targeted gene or each disrupted allele (Fig 2; S1B-G Figs). For example, when using a primer combination designed to detect intact Tb*csb* (Tbcsb-KO1/Tbcsb-q2) DNA amplification generated a band of the expected size (approximately 1.4 kb) from gDNA extracted from wild type parasites with no band(s) observed in DNA isolated from *T. brucei Tb*csb**Δ cells (Fig 2A upper panel; S1B Fig). In contrast, primer combinations that detect the hyg-(Tbcsb-KO1/hyg) or *neo-* (Tbcsb-KO1/neo) interrupted alleles generated amplicons of the predicted size (approximately 1. 7 and 1.5 kb, respectively) but only from the gDNA purified from *T. brucei Tb*csb**Δ null mutant line: No band(s) was observed when using *T. brucei* wild type gDNA as template.

**Figure 2:**
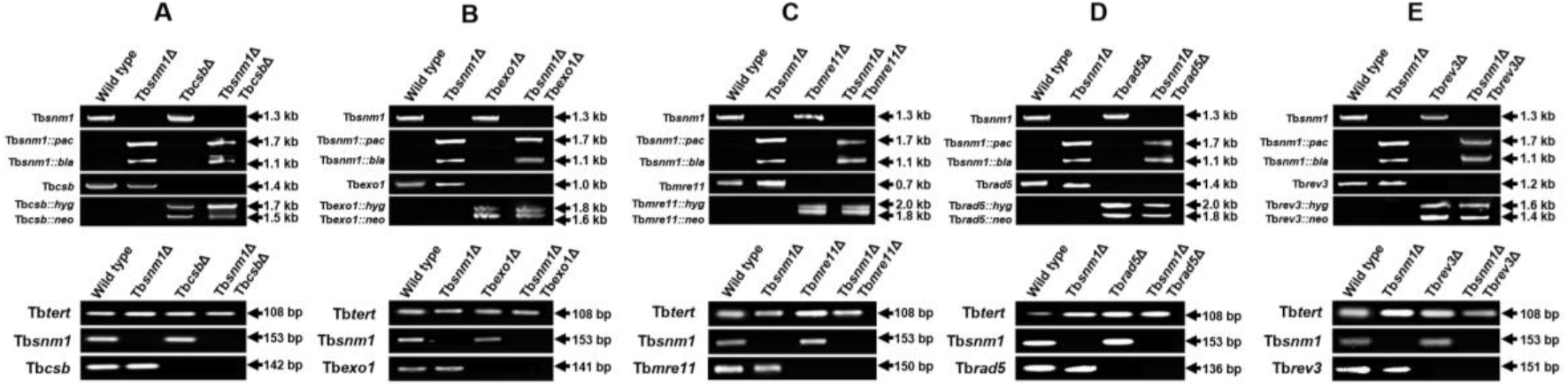
Validation of *T. brucei* null mutant lines. (A-E; upper panels). Amplicons (in kbp) corresponding to intact Tbsnm1 **(A-E)**, *Tbcsb* **(A)**, Tbexo1 **(B)**, Tb*mre11* **(C)**, Tb*rad5* **(D)** or Tb*rev3* **(E)** and their *hyg-, neo-, pac-* or bla-disrupted counterparts were generated from template genomic DNAs extracted from the various *T. brucei* lines indicated. (**A-E; lower panels**). Amplicons (in bp) corresponding to intact Tb*snm1* **(A-E)**, *Tbcsb* **(A)**, Tb*exo1* **(B)**, Tb*mre11* **(C)**, Tb*rad5* **(D)** and Tb*rev3* **(E)** were generated from template cDNAs derived from total RNA extracted from the various *T. brucei* lines indicated. The integrity of cDNAs (and hence RNAs) was evaluated by amplification of a 108 bp control fragment, Tbtert. The primer sequences and combinations used for each amplification are listed in S2 and S3 Tables, respectively.

To show that each *T. brucei* null mutant line was not expressing the targeted gene, DNA amplification reactions were performed on cDNA templates using primer combinations that generated fragments specific for the intact, targeted gene (Fig 2). For example, when using a primer combination designed to generate a Tb*csb* specific amplicon (Tbcsb-q1/Tbcsb-q2), a single band of the expected size (approximately 140 bp) was observed in cDNA synthesised from total RNA extracted from wild type parasites with no band(s) detected in material derived from *T. brucei Tb*csb**Δ (Fig 2A lower panel). To confirm that RNA had been extracted from both cell lines and that cDNA had indeed been made, control reactions amplifying Tb*tert* were conducted in parallel. For all tested samples a band of the expected size (approximately 100 bp) was observed.

The above PCR-based strategies were extended to validate *T. brucei* lines null for *Tbmre11, Tbexo1, Tbrad5* or Tbrev3 (Figs 2B-E). In each case, this confirmed that integration of the input DNA fragments had successfully occurred into the parasite genome and established that the deleted transcript was not being expressed in the appropriate null mutant line.

### Characterisation of *T. brucei* single null mutant lines

To assess whether lack of a given DNA repair activity affected parasite growth, the cumulative growth properties of null mutants was determined (S2A Fig). In most cases (Tb*exo1*Δ, Tb*csb*Δ, Tb<u>rad5</u>Δ and Tb<u>rev3</u>Δ), the lines grew at the same rate as wild type with the cultures having a doubling time of around 370 minutes. In contrast, cells lacking TbMRE11 exhibited a slight growth defect, having a mean generation time of 450 minutes, in keeping with previous observations [46, 47]. To examine these growth characteristics further, the cell cycle progression of Tb*mre11*Δ and Tb*csb*Δ cells was evaluated relative to wild type and Tb*snm1*Δ lines (S2B Fig). Asynchronous cultures of bloodstream form parasites in the exponential phase of growth were fixed and the ratio of nuclear (N) and mitochondrial (known as the kinetoplast (K)) genomes within each trypanosome determined with this providing a reliable marker for where that cell is within the cell cycle [53–55]. *T. brucei* in the G1/S phase of the cell cycle have a 1K1N arrangement, those that possess a 2K1N ratio are said to be in the G2/M phase while trypanosomes displaying a 2K2N profile are in the post M phase. For wild type *T. brucei, Tbsnm1*Δ and Tb*csb*Δ, most cells (approximately 75%) in the asynchronous population were in G1/S phase, with about 20% in the G2/M phase and around 5% in the post M phase of the cell cycle. In contrast, fewer Tb*mre11*Δ cells (approximately 55%) were in the G1/S phase, with a concomitant increase (approximately 40%) of cells in the G2/M phase of the cell cycle (S5B Fig). Therefore, the increased mean generation time exhibited by TbMRE11-deficient parasites is mostly due to a delay at the G2/M phase of the cell cycle. In other organisms, MRE11 as part of the MRN complex, functions to recognise DSBs generated during DNA replication then aids in recruitment of ATM to the site of DNA damage. Perturbation of this sensor mechanism effects downstream ATM-dependent processes such as progression through the G2/M checkpoint [56].

To evaluate whether deletion of Tb*csb*, Tb*mre11*, Tb*exo1*, Tb<u>rad5</u> or Tbrev3 from the *T. brucei* genome altered the parasites sensitivity to DNA damage, the null mutants were grown in the presence of hydroxyurea, phleomycin or methyl methanesulfonate, or cultured following exposure to a single dose of UV, and the EC_50_ values towards each treatment determined (Table 1; S2C & D Figs). Cells lacking TbCSB were 5-fold more susceptible to UV than wild type, in keeping with previous observations [41], while parasites lacking TbEXO1 displayed an increased sensitivity (2-fold) towards phleomycin. In keeping with the published literature, TbMRE11 deficient parasites were more susceptible to phleomycin (12-fold) and methyl methanesulfonate (2.5-fold), agents that promote DSBs (Table 1: S2C & D Figs) [46, 47]. Intriguingly, these mutants were also shown to be 4.9-fold more sensitive to UV, a trait not previously reported, with this phenotype potentially stemming from DSB formation caused by the arrest of replication forks at UV-induced damaged sites [57]. Cells lacking TbRAD5 and TbREV3 were as sensitive to all the DNA damaging treatments tested as wild type indicating that these do not play a front line role in resolving the lesions generated by the tested treatments.

**Table 1.**
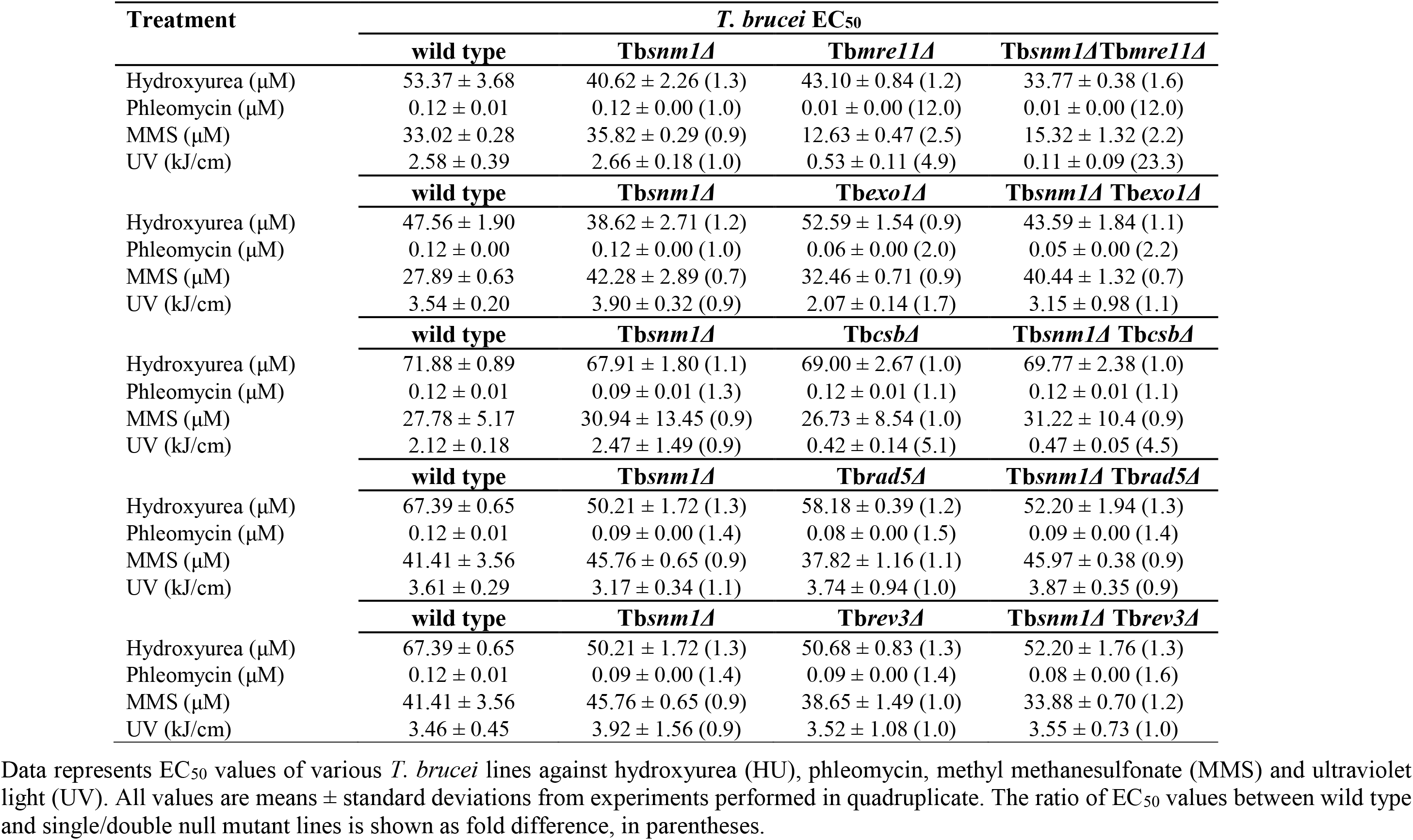
Susceptibility of *T. brucei* null lines to DNA damaging treatments.

### Susceptibility of *T. brucei* single null mutants towards mechlorethamine

The above growth inhibition assays were extended to investigate the phenotype displayed by *T. brucei* null mutants towards mechlorethamine, an archetypal ICL inducing agent. The resultant data was plotted as dose response curves from which EC_50_ values were extrapolated (Fig 3). Previous work has shown that TbSNM1 plays an important role in resolving the damage caused by mechlorethamine with our data confirming this earlier finding [42]: Cells lacking TbSNM1 are >8-fold more susceptible to this compound as compared to controls. A similar alteration in sensitivity was also observed in cells deficient in TbMRE11, TbCSB or TbEXO1 although the difference in EC_50_ values displayed by these 3 mutants relative to wild type was not as great as that noted for Tb*snm1*Δ cells; Tb*mre11*Δ, Tb*csb*Δ and Tb*exo1*Δ cells were 3-, 4- and 2-fold more susceptible to mechlorethamine than wild type, respectively (Figs 3A-C) This could suggest the relative importance of each enzyme in *T. brucei*’s so-called ICL REPAIRtoire. In contrast, cells lacking TbRAD5 or TbREV3 displayed wild type sensitivities towards the ICL inducing agent indicating they play no significant role in resolving this type of damage (Figs 3D-E). To confirm that the above phenotyping was due to the engineered gene deletion/disruption events and not due to off target effects, the susceptibility of all mutants towards the non-DNA damaging trypanocidal agent difluoromethylornithine was analysed (S3 Fig). In all cases the null lines were as equally sensitive to this compound as each other and the wild type, providing evidence that the above susceptibility profiles are specific for each DNA damaging treatment.

**Figure 3:**
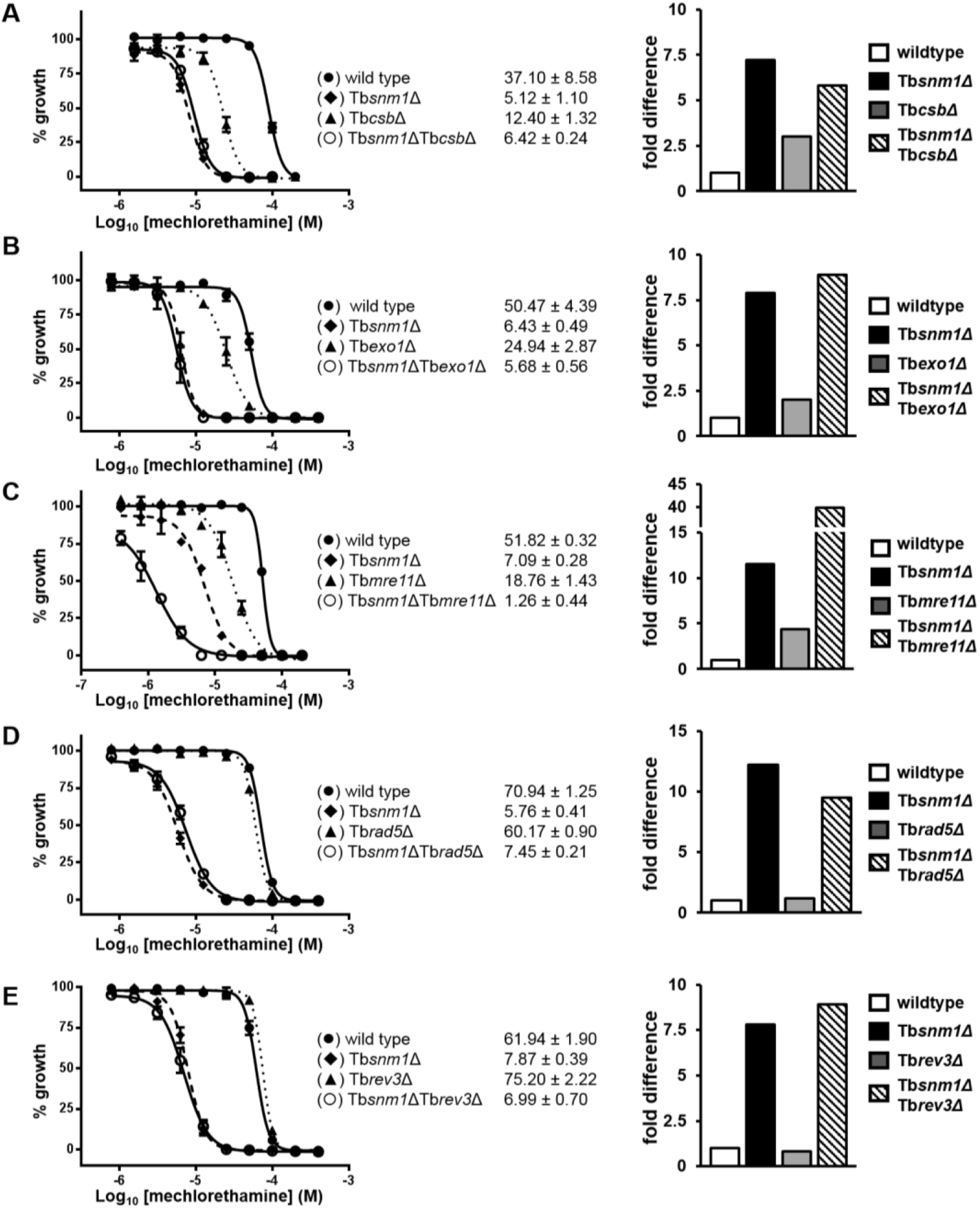
Susceptibility of *T. brucei* null mutants towards mechlorethamine. Left hand panels: Dose response curves and EC_50_ values (in μM) of *T. brucei* lines towards mechlorethamine. All data points are mean values ± standard deviations from experiments performed in quadruplicate. Right hand panels: The susceptibility of the *T. brucei* single and double null mutant lines against mechlorethamine, as judged by their EC_50_ values, were compared and expressed as a fold difference relative to wild type.

As TbMRE11 functions in *T. brucei*’s HR pathway, the above studies were extended to evaluate whether parasites deficient in other components of this repair system (TbBRCA2 or TbRAD51) exhibited altered susceptibilities towards mechlorethamine: TbBRCA2 and TbRAD51 correspond to two known FA proteins, FANCD1 and FANCR, respectively (Fig 1D). From the resultant dose response curves and EC_50_ values (Fig S4), parasites lacking TbRAD51 or TbBRCA2 were shown to be 6- or 3-fold more sensitive to the ICL inducing agent than controls, respectively, thus implicating these activities in the trypanosomal ICL repair network. The above data implicates TbSNM1, TbCSB, TbEXO1, TbMRE11, TbRAD51 and TbBRCA2 in the bloodstream form *T. brucei* ICL repair system and thus constitute a part of this parasite’s ICL REPAIRtoire.

### Mechlorethamine-induced γH2A formation in *T. brucei* single null mutants

Nuclear genome damage in many eukaryotes including *T. brucei* often leads to phosphorylation of H2A to form γH2A an alteration frequently used to monitor for DNA damage including DSB formation [55, 58, 59]. To determine if mechlorethamine can promote this post-translational modification in *T. brucei*, protein extracts generated from wild type parasites continuously grown in the presence of mechlorethamine (30, 10 and 3 μM) were probed for γH2A formation using an antiserum against this modified histone (Fig 4A), with the band intensities normalised against untreated controls (Fig 4B): A similar analysis was performed on the same extracts using enolase as loading & normalisation control. In wild type cells and at all mechlorethamine concentrations tested, an increase in the γH2A signal was observed over the first 4 hours of treatment with the marker intensity then declining over the next 4 hours indicating resolution of the DNA damage. When these studies were extended to the null lines, various outcomes were observed. TbSNM1- and TbEXO1-deficient cells behaved similarly to wild type cells. In Tb*csb*Δ cells, the formation of γH2A was delayed, and continued to increase throughout the experiment suggesting that these parasites may be unable to effectively recognise and resolve any ICL-induced DSBs. Intriguingly, when following γH2A formation in cells lacking TbMREll, no alteration in signal intensity was noted at any time point across any of the mechlorethamine concentrations used indicating that this enzyme plays an important role in phosphorylation of histone H2A in response to DNA damage [60–62].

**Figure 4:**
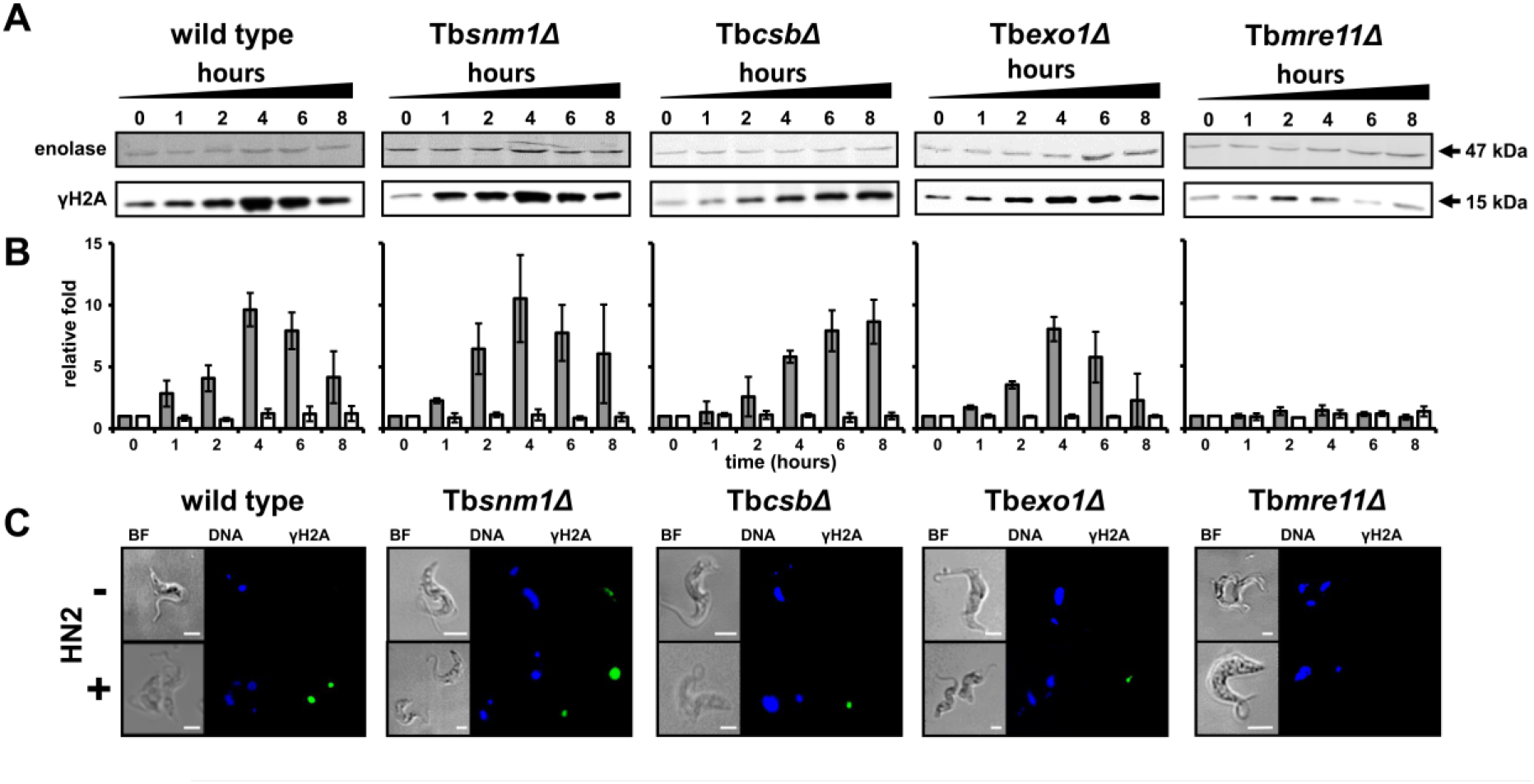
γH2A formation in mechlorethamine treated *T. brucei*. **(A)**. *T. brucei* wild type, Tb*snm1*Δ, Tb*csb*Δ, Tb*exo1*Δ and Tb*mre11*Δ were treated with 30 μM mechlorethamine. Cell lysates were generated at time intervals (0, 1, 2, 4, 6, and 8 hours) and analysed by western blot using anti-T. *brucei* enolase (loading control) and anti-T. *brucei* γH2A antiserum. Treatment with other mechlorethamine concentrations (10 and 3 μM) resulted in similar γH2A responses (data not shown). **(B)**. The γH2A and enolase signal intensities obtained from western blots containing extracts from three independent mechlorethamine treated cultures were determined using Image Studio™ Lite (Li-COR Biosciences) and normalised against untreated controls. The data is expressed as a mean ± standard deviation relative fold difference. **(C)**. The pattern of DNA (blue) and γH2A (green) staining in *T. brucei* treated (+) for 4 hours with mechlorethamine (HN2) was compared with signals observed in untreated (-) cells. The cells were examined by florescence microscopy and the brightfield (BF) image captured. Scale bar = 5 μM.

To further evaluate mechlorethamine and DNA damage in mutant lines, cells treated for 4 hours in the presence of the ICL-inducing agent (30 μM) were analysed using immunofluorescence microscopy to detect γH2A (Fig 4C). For wild type, Tb*snm1*Δ, Tb*csb*Δ and Tb*exo1*Δ, a discrete signal was observed within the nucleus showing that DNA damage was taking place. This pattern also was detected in wild type, Tb*snm1*Δ and Tb*exo1*Δ cells that had been treated with mechlorethamine for 2 hours but not in Tb*csb*Δ parasites. In contrast, and confirming the western blot data, no signal was observed in mechlorethamine treated TbMRE11 null parasites.

### Assessing the interplay between trypanosomal ICL repair enzymes

To evaluate the functional relationship between components of the trypanosomal ICL repair system, the genes encoding for TbEXO1, TbMRE11 or TbCSB were disrupted in a *Tbsnm1-* deficient *T. brucei* line to create a series of double null mutant lines. Additionally, TbRAD5 and TbREV3 although not primarily involved in resolving ICLs, were also taken forward to determine whether their activities in this repair network becomes apparent in the absence of TbSNM1: The role of some yeast factors in ICL repair are only apparent when the activity of another DNA repair protein is missing [63]. To generate the double null mutant lines, the hyg-and *neo*-based integration vectors for each targeted gene were linearised and sequentially transformed into *T. brucei Tb*snm1**Δ cells (in this mutant line, Tbsnm1 has been disrupted using vectors based around resistance cassettes that includes the gene encoding for blasticidin-S-deaminase *(bla)* or puromycin N-acetyltransferase (*pac*) [42]). Following selection, all putative double null clones were validated using the PCR-based strategies outlined previously with this confirming that the introduced DNA fragments had integrated into the *T. brucei* genome and that the recombinant parasites were no longer expressing both disrupted genes (Fig 2).

Analysis of the double null mutants revealed that parasites lacking TbSNM1 and TbCSB, TbEXO1, TbRAD5 or TbREV3 grew roughly at the same rate as wild type while cells deficient in TbSNM1 and TbMRE11 exhibited a slightly longer mean generation time comparable to that noted for the *T. brucei Tb*mre11**Δ line. Additionally, phenotypic screening with DNA damaging treatments demonstrated that the susceptibility displayed by the double null mutant lines were generally equivalent to the more sensitive phenotype shown by the single null for each gene pairing (Table 1). For example, all Tb*csb*-deficient cell types were approximately 5fold more susceptible to UV than wild type or Tb*snm1*Δ such that the *Tbsnm1 ΔTb*csb**Δ line exhibited an EC_50_ similar to Tb*csb*Δ parasites. As such, lack of TbSNM1 and TbCSB activities in the same cell does not lead to an increase in susceptibility to UV. The only situation where a combinatorial effect on susceptibility was noted related to cells lacking TbSNM1 and TbMRE11. Intriguingly, such double null mutant parasites were more than 20-fold more sensitive to UV-induced lesions relative to *T. brucei* wild type and Tb*snm1*Δ, and 4-fold more sensitive relative to Tb*mre11*Δ (Table 1). This difference could reflect that TbSNM1 does play a secondary role in resolving UV-induced damage, a phenotype that is only apparent in cells compromised in the HR pathway.

The phenotypic screens were extended to investigate the susceptibility that *T. brucei* double null mutants display towards mechlorethamine with the resultant data plotted as dose response curves from which EC_50_ values were extrapolated (Fig 3). This revealed that cells lacking TbSNM1 and TbMRE11 were hypersensitive towards this ICL-inducing agent relative to wild type and the Tb*snm1*Δ or Tb*mre11*Δ lines showing that these two nucleases function in a non-epistatic fashion and do not operate in the same ICL repair system (Fig 3C). In contrast, parasites deficient in TbSNM1 and TbCSB or in TbSNM1 and TbEXO1 exhibit dose response sensitivities to mechlorethamine similar to Tb*snm1*Δ parasites, with the double null lines being more susceptible to this compound than wild type and Tb*csb*Δ or Tb*exo1*Δ cells (Figs 3A & B). The type of patterns observed indicate that TbSNM1 functions epistatically with TbCSB and TbEXO1. When the susceptibility phenotype of trypanosomes lacking TbSNM1 and TbRAD5 or TbREV3 towards mechlorethamine was assessed, dose response curves similar to that obtained using control lines were observed (Figs 3D & E). This suggests that TbRAD5 and TbREV3 do not play a role in the *T. brucei* ICL repair networks.

To further investigate any epistatic/non-epistatic relationships, double null lines lacking TbCSB and TbMRE11, TbEXO1 and TbMRE11 or TbCSB and TbEXO1 activities were made and the resultant lines validated (Figs 5A-C). Evaluation of dose response curves of parasites lacking TbCSB and TbEXO1 towards mechlorethamine revealed that these cells exhibited a susceptibility pattern similar to Tb*csb*Δ trypanosomes, with the double null lines being more sensitive to this treatment than wild type and Tb*exo1*Δ cells (Fig 5D). This clearly demonstrates that these *Tbcsb* is epistatic with respect to Tbexo1 in the *T. brucei* ICL repair system. In contrast, *Tb*csb*ΔTb*mre11**Δ and Tb<u>exo1</u> ΔTb*mre11*Δ trypanosomes were hypersensitive to this ICL-inducing agent relative to wild type cells and the corresponding single null mutant lines providing evidence that TbCSB and TbEXO1 both function in a non-epistatic fashion with respect to TbMRE11 and as such the former two activities operate in an ICL repair system distinct from that the one the latter is involved in (Figs 5E and F).

**Figure 5:**
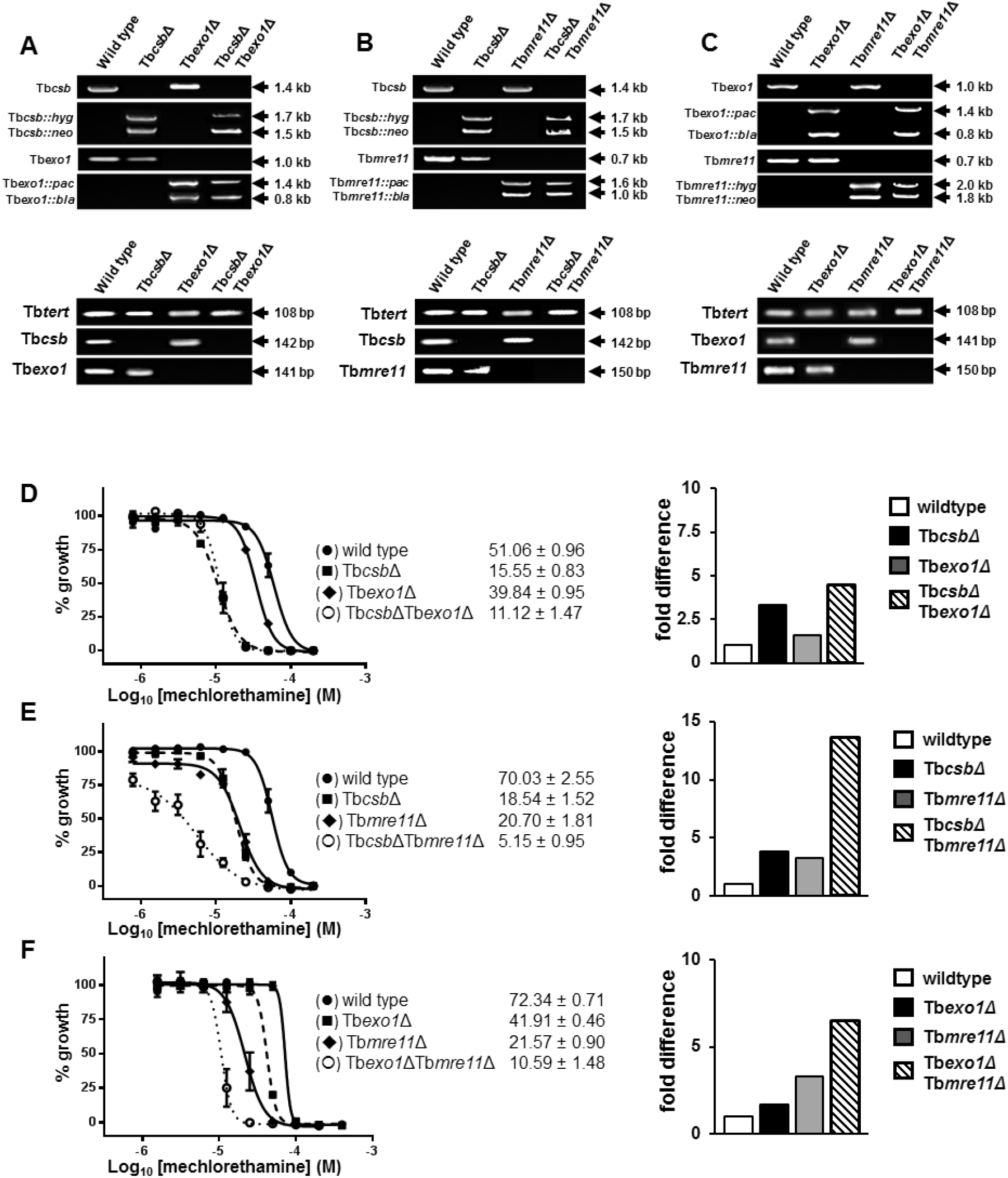
Evaluating the interplay between TbCSB, TbMRE11 and TbEXO1 in the *T. brucei* ICL repair system. (A-C; upper panels). Amplicons (in kbp) corresponding to *Tbcsb* **(A & B)**, Tb*exo1* **(A & C)** or Tb*mre11* **(B & C)** and their *hyg-, neo-, pac-* or bla-disrupted counterparts were generated from template genomic DNAs extracted from the various *T. brucei* lines indicated. (**A-C; lower panels**). Amplicons (in bp) corresponding to intact Tb*csb* **(A & B)**, Tb*exo1* **(A & C)** or *Tbmre11* **(B & C)** were generated from template cDNAs derived from total RNA extracted from the various *T. brucei* lines indicated. The integrity of cDNAs (and hence RNAs) was evaluated by amplification of a 108 bp control fragment, Tbtert. The primer sequences and combinations used in for each amplification are listed in S2 and S3 Tables, respectively. **(D-F)**. Dose response curves and EC_50_ values (in μM) of *T. brucei* lines towards mechlorethamine are shown. All data points are mean values ± standard deviations from experiments performed in quadruplicate. The susceptibility of the *T. brucei* single and double null mutant lines against mechlorethamine, as judged by their EC_50_ values, were compared and expressed as a fold difference relative to wild type.

All of the double null mutant data clearly shows that *T. brucei* expresses at least two distinct ICL repair systems with one involving the concerted action of TbSNM1, TbCSB and TbEXO1 with the other using TbMRE11. It is tempting to speculate that given TbCSB plays a role in TC-NER and that the former SNM1-dependent pathway functions to resolve ICLs encountered during DNA transcription while the MRE11-dependent system helps remove ICLs encountered during DNA replication.

## Discussion

Throughout its cell and life cycle *T. brucei* may be exposed to a range of endogenous metabolites and environmental stimuli that in other organisms result in ICL formation [64–66]. Despite ICLs being highly toxic, very little is known about the mechanisms *T. brucei* employs to resolve such lesions. The ICL repair networks expressed in yeast and mammalian cells can be distinguished by how these systems are activated and the factors involved [18, 21, 22, 24, 67]. Throughout the cell cycle, but most prominently during the G1/G0-phases, replication-independent systems operate with these mechanisms triggered by global genome surveillance processes (by GG-NER) or stalling of transcription complexes (by TC-NER) [25, 26, 68]. Following lesion detection, networks are invoked that involve components from the NER and/or MMR pathways which together unhook then resect the ICL from the genome with any excised genetic material replaced by TLS [28, 69]. In contrast, during the S-phase of the cell cycle, replication-dependent mechanism(s) function to eliminate ICLs [31, 35–37, 63, 70]. In such networks, recognition of an ICL stalled DNA replication fork(s) occurs through components of the FA or Fanconi-like systems that firstly cause reversal of the replication fork(s) then guide predominantly HR effector proteins to the site of damage. In conjunction with the endonucleases centred on XPF-ERCC1 activity, these cooperate to remove and replace the cross linkage from the genome using homologous sequences as a repair template. Here, using comparative genomics in conjunction with classical genetic approaches we have identified several putative trypanosomal ICL repair proteins and established whether the *in silico* assignments were of biological relevance. This revealed that enzymes from the *T. brucei* NER, HR and/or MMR pathways contribute to this organism’s ICL REPAIRtoire to resolve damage generated by mechlorethamine and that these activities function across two distinct networks.

An informatics approach was employed to determine the extent of the *T. brucei* ICL REPAIRtoire. Using protein sequences from representative members of the yeast and/or mammalian ICL repair systems as bait, genome database searches were performed identifying trypanosomal homologues of the TC-NER helicase CSB, the MMR/HR exonuclease EXO1 and the REV3 catalytic subunit of pol *ζ* involved in error-prone TLS were identified. Additionally the ICL repair specific nuclease TbSNM1, several HR components including MRE11, RAD51 and BRCA2, and RAD5, a DNA helicase/ubiquitin ligase that can promote HR in a template switching, error-free DNA damage tolerance pathway, were found. Intriguingly, no discernible homologues for any component of the FA core complex (FANCA, −B, −C, −E, −F, −G, −L and −T plus the ancillary factors FAAP20, −24 and −100), recruitment factors (FANCD2A and −I) or FA-like proteins (Mph1 and Slx4) were identified suggesting the ICL-mediated stalling of DNA replication forks is recognised by trypanosomal mechanisms distinct from those typically expressed by other eukaryotes with this potentially triggering unusual downstream responses.

In many cases, the use of null mutant lines has provided the primary route for the identification of which DNA repair factors function in an organism’s ICL REPAIRtoire and in deciphering how these components interact [45, 63, 71, 72]. Transferring this approach to *T. brucei*, lines lacking a single potential ICL repair factor were made (TbCSB, TbEXO1, TbMRE11, TbRAD5 or TbREV3) or sourced (TbSNM1, TbRAD51 or TbBRCA2 [42, 50, 52]) with these subsequently phenotyped using mechlorethamine as a selective agent. For most parasite lines tested, an increased susceptibility to the ICL-inducing agent was observed, demonstrating that TbSNM1, TbCSB, TbEXO1, TbMRE11, TbRAD51 and TbBRCA2 all contribute to the trypanosomal ICL REPAIRtoire. Comparison of the degree of susceptibility of each mutant in relation to controls suggests that TbSNM1 may play a prominent role in resolving crosslinks while TbCSB, TbMRE11, TbRAD51 and TbBRCA2 all represent important factors in these repair systems: Tb*snm1*Δ parasites exhibited the highest fold (around 7.5-fold) difference in susceptibility while Tb*csb*Δ, Tb*mre11*Δ, Tb*rad51*Δ and Tb*brac2*Δ cells displayed moderate (between 3- to 5-fold) changes. In the case of TbEXO1, cells lacking this exonuclease displayed about a 2-fold increase in sensitivity indicting that this enzyme may play an ancillary role in the trypanosomal ICL repair network. When these screens were extended to TbRAD5- or TbREV3-deficient parasites no obvious differences in susceptibility was observed. As with other organisms, trypanosomes express several error-free and error-prone DNA damage tolerance mechanisms and it is plausible that these other systems may complement for the lack of TbRAD5 or TbREV3 activity in the corresponding *T. brucei* null line [73].

Resolution of ICLs involves the concerted action of various enzymes from different ‘classical’ DNA repair pathways operating across several cell cycle dependent networks and all within a single cell. In yeast, the interplay of ICL repair factors and their assignment to a given pathway frequently involves use of fungal lines lacking multiple DNA repair activities and determining their susceptibility towards an ICL-inducing agent [45, 63, 71, 72]. Such studies can then help inform the situation in other organisms. To evaluate the interactions of TbSNM1, TbCSB, TbEXO1 and TbMRE11, *T. brucei* lines lacking various combinations of two of these activities were generated with the parasites subsequently phenotyped against mechlorethamine. This revealed that TbSNM1, TbCSB and TbEXO1 function in one *T. brucei* ICL repair system with TbMRE11 operating as part of a second distinct network. This situation resembles that noted in other organisms although there are some key differences. In *S. cerevisiae*, PSO2 (the yeast SNM1 homologue) displays a non-epistatic interaction with EXO1 and all tested DSB repair protein, including MRE11 [45, 72, 74, 75]. It also associates with several NER factors such as RAD1 (XPF), RAD3 (XPD), RAD4 (XPC) and RAD14 (XPA) [22, 72] although whether it interacts with RAD26 (the yeast CSB homologue) remains unclear [22]. In humans, the role played by SNM1 is reportedly more complex partly because each cell can express multiple isoforms of this family of nuclease [76]. In the case of SNM1A, the human homologue most like yeast PSO2, a direct interaction with CSB has been established [26] with associations also noted for other NER factors such as XPC, DDB2 and XPF–ERCC1 [27, 68]. In a situation contrary to that observed in yeast and *T. brucei*, SNM1A also appears to play a role in replication-dependent ICL resolving mechanisms as it reportedly co-localises with the HR factors such as MRE11 and BRCA1 while human cells with depleted levels for this activity exhibit an increase in replication associated DSBs [27, 77]. Additionally, human cells also express a second SNM1/PSO2 isoform called SNM1B that has been shown to physically interact with MRE11 *via* an MRN binding site with this activity facilitating direct DNA repair at the site of stalled DNA replication forks, a HR-dependent process that occurs during S-phase of the cell cycle [76, 78, 79]. Therefore, it appears that in single-celled eukaryotes members of the SNM1/PSO2 family function only in replication-independent ICL repair systems while in higher eukaryotes this class of nuclease additionally play roles in replication-dependent ICL repair processes.

Based on the pathways reported in other organisms coupled with observations made by ourselves and by Machado *et al* [41], we postulate that in bloodstream form *T. brucei* ICLs encountered throughout the cell cycle as a result of global genome surveillance through GG-NER (*e.g*. TbXPC, TbDDB, TbRAD23) or transcription complex stalling by TC-NER (*e.g*. TbCSB) recognition mechanisms trigger TbXPF-TbERCC1 (and possibly XPG) incision of one DNA strand at sites flanking the ICL (Fig 6A). The resultant unhooked sequence subsequently undergoes nucleolytic processing, a process carried out by TbSNM1 (and possibly TbEXO1), up to and beyond the crosslink, leaving a single nucleotide tethered to the complementary strand. Replication Protein A binds to the single stranded gap generated by the unhooking activity that *via* activated PCNA recruits TLS DNA polymerases to this site. These in conjunction with DNA ligases restore the integrity of the DNA strand. To completely remove the crosslinked nucleotide from the other DNA strand a second round of incision followed by TLS activity occurs to generate the undamaged dsDNA structure. In contrast and representing a DNA replication-dependent mechanism, two replication forks converge upon an ICL to form an X-shaped structure (Fig 6B). Following recognition by an unknown mechanism that does not involve the FA core or FA-like proteins, incision of one DNA strand at sites flanking the ICL occurs possibly through the action of TbXPF-TbERCC1, TbMUS18 and/or TbFAN1. This results in unhooking of the ICL from one DNA strand, formation of a single stranded gap and generation of DSBs, with the latter detected through the TbMRE11-mediated formation of γH2A. The single stranded gap can be filled as described above through TLS DNA polymerase/DNA ligase activities with the unhooked sequence removed and replaced by a second round of incision and TLS: The unhooked sequence may undergo nucleolytic processing prior to the 2^nd^ incision event in an SNM1-independent mechanism possibly involving TbEXO1. The DSBs can then be repaired by HR involving TbRAD51 and TbBRCA2, using the newly formed dsDNA as a template. The two proposed trypanosomal pathways both employ TLS activities to fill single strand gaps that are associated with unhooking events. However certain DNA polymerases such as pol *ζ* are error prone and as such their activity may introduce base mismatches in the newly ‘repaired’ sequence. To overcome such secondary lesions, MMR mechanisms possibly involving TbEXO1 can operate although these may create point mutations relative to the initial sequence.

**Figure 6:**
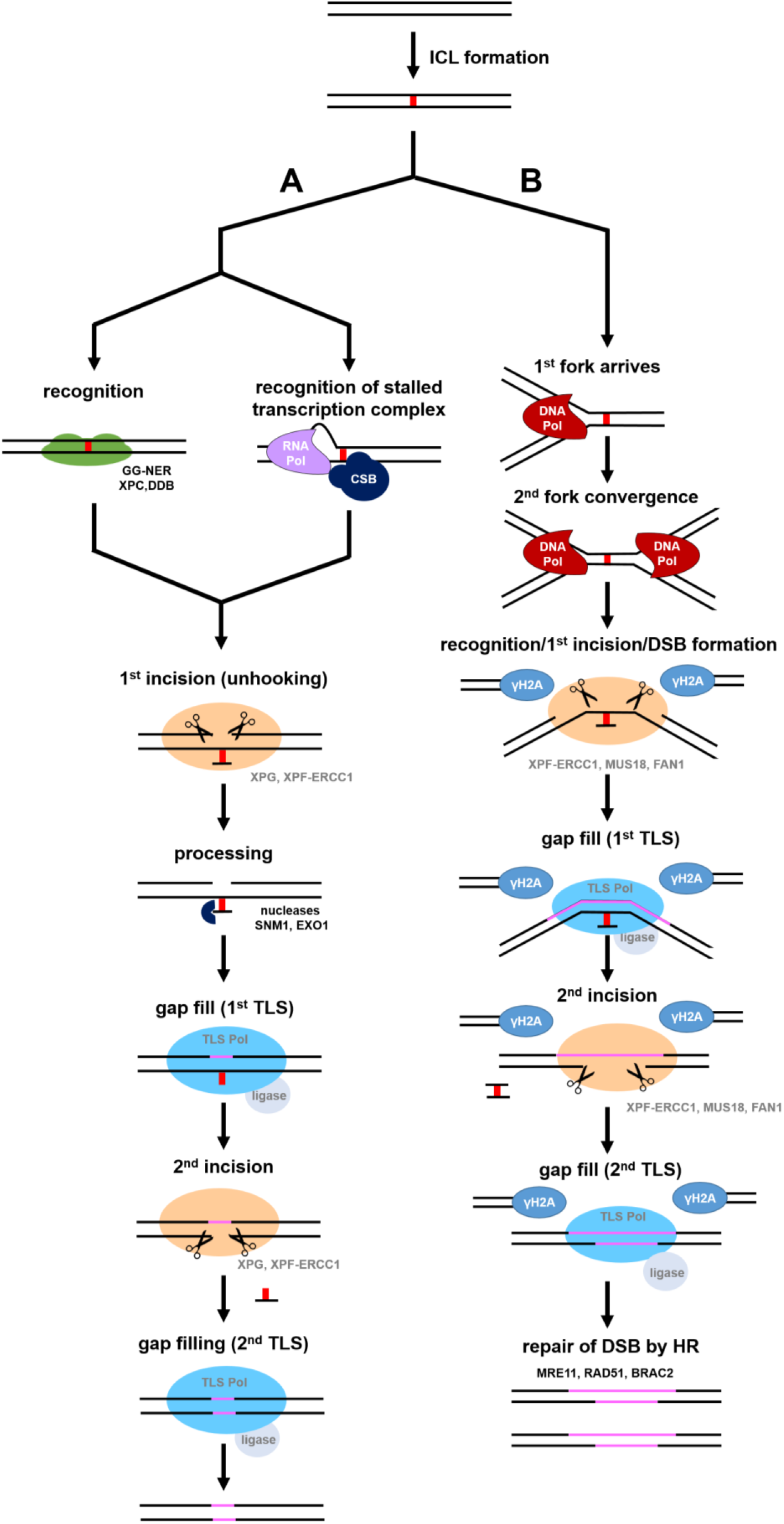
Overview of the *T. brucei* ICL REPAIRtoire. In the *T. brucei* DNA replication-independent ICL repair network **(A)**, ICLs (red bar) recognised as a result of global genome surveillance (TbXPC & TbDDB of GG-NER [41] or stalling of transcription complexes (TbCSB of TC-NER) triggers NER-mediated incision of the DNA backbone, a process that in other organisms involves XPF-ERCC1. This generates an ‘unhooked’ crosslinked oligonucleotide that undergoes nucleolytic processing (TbSNM1, TbEXO1) resulting in a single nucleotide tethered to the complementary strand, and a single stranded gap. Replication protein A binds to the single stranded DNA gap, promotes activation of PCNA that in turn recruits error free or error prone TLS DNA polymerases which synthesise new DNA (in purple) opposite the unhooked lesion, with the DNA backbone re-joined by DNA ligase: As trypanosomes express multiple TLS DNA polymerases the role of one enzyme in ICL repair (e.g. TbREV3) could be complemented by other related activities. A second round of NER-mediated incision of the DNA backbone at sites flanking the ‘unhooked’ crosslinked single nucleotide followed by TLS results in complete removal of the covalent linkage and restoration of dsDNA. In the *T. brucei* DNA replication-dependent ICL repair network **(B)**, two DNA replication forks converge upon an ICL, causing collapse of each complex. Following recognition (trypanosomal mechanism unknown), incision of the DNA backbone occurs, possibly through the activity of XPF-ERCC1, MUS18 or FAN1 endonucleases, resulting in double strand DNA break (DSB) formation (detected by TbMRE11-mediated formation of γH2A), unhooking of the ICL and generation of a single stranded gap: The unhooked ICL may undergo nucleolytic processing (TbEXO1). A first round of TLS activity results in filling of the single stranded gap associated with formation of the ‘unhooked’ sequence followed by a second round of NER-mediated incision that removes the ICL from the genome, and TLS that restores the dsDNA structure. The DSBs are then repaired by HR (involving TbRAD51 and TbBRCA2) using the newly formed dsDNA as template. Activities that have been shown to be involved in ICL repair are highlighted in black (XPC, DDB1, SNM1, EXO1, MRE11, RAD51 and BRCA2) or white (CSB) whereas activities that are hypothesised to function in these systems (XPF-ERCC1, XPG, MUS81, FAN1, TLS Pols & ligase) are in grey.

We have now shown that *T. brucei* expresses several enzymes from a number of classical DNA repair pathways that also function to resolve ICLs. As in other organisms, these activities operate across multiple, distinct pathways that we postulate function to fix lesions encountered at different points in the cell cycle. The comparative analysis performed here on a relatively small number of targets has already shown that the ICL resolving mechanisms employed by *T. brucei* share similarities to those expressed by other eukaryotes but has also highlighted some intriguing differences. By further unravelling the ICL repair systems additional variations may be identified. Such disparities between the trypanosomal and mammalian networks could identify potential targets that may be exploitable in terms of drug development such that the specific inhibition of a parasite ICL repair factor(s) may render the pathogen more susceptible to compounds that promote ICL formation.

## Material and methods

### Parasite culture

Bloodstream form *Trypanosoma brucei brucei* MITat 427 (clone 221a) (designated as wild type) was grown in HMI-9 (Invitrogen) medium supplemented with 3 g l^−1^ sodium bicarbonate, 0. 014 % (v/v) β-mercaptoethanol and 10 % (v/v) foetal bovine serum (FBS) at 37 °C under a 5 % (v/v) CO2 atmosphere [80]. Derivatives of the ‘221’ line null for DNA repair enzymes (S1 Table), including cells lacking Tbsnm1, *Tbrad51* and *Tbbrca2* [42, 50, 52], were maintained in the above growth medium containing 2.5 μg ml^−1^ hygromycin, 10 μg ml^−1^ blasticidin, 2 μg ml^−1^ puromycin and/or 2 μg ml^−1^ G418.

### Chemicals & treatments

Agents that promote DNA damage were obtained from Sigma-Aldrich (methyl methanesulfonate, hydroxyurea, phleomycin) or Cambridge Biosciences (mechlorethamine) while UV irradiation was performed with a Stratalinker UV crosslinker (Stratagene). The trypanocidal selective compounds blasticidin, puromycin, G418 and hygromycin were sourced from Melford Laboratories Ltd with difluoromethylornithine supplied by Prof. Mike Barrett (University of Glasgow).

### Nucleic acids

The Tbcsb (Tb927.7.4080), Tbmre11 (Tb927.2.4390), Tbexo1 (Tb927.8.3220), *Tbrad5* (Tb927.7.1090) and Tbrev3 (Tb927.8.3290) open reading frames and flanking regions were identified following reciprocal BLAST searches of TriTrypDB (http://tritrypdb.org/tritrypdb/) and NCBI database (https://www.ncbi.nlm.nih.gov/) using the human and/or yeast orthologue as bait, and analysis of trypanosomal sequence information in the published literature [41, 46, 47, 81]. Deduced protein sequences were aligned to the counterparts expressed by other organisms using CLUSTALΩ (http://www.ebi.ac.uk/Tools/msa/clustalo/), domain structures evaluated using HMMR (http://www.ebi.ac.uk/Tools/hmmer/) and regions of homology highlighted using BoxShade (https://embnet.vital-it.ch/software/BOX_form.html).

### Construction of null mutant lines

The vectors used to delete or interrupt a target gene in the *T. brucei* genome were generated as follows. For Tbcsb, sequences corresponding to 5’ and 3’ flanking regions were amplified from *T. brucei* genomic DNA and sequentially cloned either side of a resistance cassette that includes the gene encoding for hygromycin B phosphotransferase or neomycin phosphotransferase. The vectors designed to disrupt Tbexo1, *Tbmre11, Tbrad5* or Tbrev3 were made in a similar way except that the amplified fragments were derived from the 5’ and 3’ ends of the coding sequence and these were cloned either side of a resistance cassette that included the gene encoding for *hyg, neo, bla* or pac: The primers and primer combinations used to generate the 5’ and 3’ sequences and the sizes of the amplified fragments are shown in S2 and S3 Tables, and S1A Fig. Constructs were linearized (SacI/KpnI for the hyg, *neo* and *pac* constructs and SacII/ KpnI for the *bla* vector) then transformed into bloodstream form *T. brucei* using the Human T-cell nucleofection^®^ kit and an Amaxa^®^ Nucleofector™ (Lonza AG) set to program X-001. As the *T. brucei* genes targeted here are single copy in a diploid genome, two rounds of nucleofection were needed to firstly create heterozygous and then null mutant lines. Integration of the Tbcsb-based DNA constructs into the *T. brucei* genome resulted in deletion of the entire open reading frame. In contrast, integration of the Tbexo1-, Tbmre11-, *Tbrad5-* or Tbrev3-based gene disruption constructs into the *T. brucei* genome resulted in deletion of 50, 37, 67 or 80 % of the corresponding open reading frame, respectively. The regions removed from the genome sequences are postulated to be needed for DNA binding (TbEXO1, TbMRE11, TbRAD5), nuclease (TbEXO1, TbMRE11, TbREV3), ubiquitin ligase (TbRAD5) or DNA polymerase (TbREV3) activities.

### Validation of recombinant parasites

To demonstrate that integration of a gene interruption cassette had occurred at the correct genetic loci and confirm that the *T. brucei* null mutant line was no longer expressing the targeted gene(s), DNA amplification reactions were performed on parasite genomic DNA or cDNA templates. For gene interruption constructs, the primer combinations used generated amplicons specific for the intact targeted gene or the *hyg*-, *neo*-, *pac*- and/or *bla*-disrupted allele. For all DNA’s analysed, additional reactions aimed at detecting *Tbtert* were performed on cDNA to check the integrity of these templates and as loading control. The primer sequences and combinations used in cell line validation are listed in S2 and S3 Tables.

### Growth curves

Bloodstream form *T. brucei* in the logarithmic phase of growth were seeded at 1 × 10^4^ parasites ml^−1^ in parasite growth medium and incubated at 37 °C under a 5 % (v/v) CO2 atmosphere. Each day, the cell density of each culture was measured using a Neubauer haemocytometer. When the number of parasites reached approximately 1 × 10^6^ ml^−1^, a new culture seeded at 1 × 10^4^ parasites ml^−1^ was set up. This analysis was carried out over an 8 to 14 day period. Growth curves were generated using GraphPad Prism (GraphPad Software Inc.). All growth assays were performed in triplicate and each count at each time point expressed as a mean ± standard deviation. Mean generation times were calculated using http://www.doubling-time.com/compute.php.

### Microscopy and protein analysis

Bloodstream form *T. brucei* were fixed in growth medium with an equal volume of 2 % (w/v) paraformaldehyde/PBS, washed once in phosphate buffered saline (PBS) and aliquots containing approximately 10^5^ cells air dried onto a microscope slide. For cell cycle arrest assays, the genomes of paraformaldehyde-fixed trypanosomes were stained with 4’,6-diamidino-2-phenylindole (DAPI) and trypanosomes visualized using a Leica DMRA2 epifluorescent microscope in conjunction with a C4742-95 digital camera (Hamamatsu Photonics). For immunofluorescence studies, paraformaldehyde-fixed trypanosomes were permeabilized for 15 minutes in 0.5 % (v/v) Triton-X (Sigma-Aldrich) in PBS, blocked with 50 % (v/v) FBS in PBS then labelled with rabbit anti-γH2A (Dr Richard McCulloch, University of Glasgow) antisera diluted 1:250 in 3 % (v/v) FBS in PBS. After 45 minutes, the slides were washed extensively in PBS then incubated for 45 minutes with Alexa-Fluor 488 goat antimouse (Molecular Probes) diluted 1:1000 in 3 % (v/v) FBS in PBS. Following further PBS washes, the parasite DNA was stained using Vectashield Mounting Medium containing DAPI (Vectorshield Laboratories). Images were captured using a Leica SP5 confocal microscope (Leica Microsystems (UK) Ltd). All images were processed and corrected fluorescence intensities determined using ImageJ.

For western blot analysis, protein extracts from approximately 2 x 10^6^ trypanosomes were probed with rabbit anti-γH2A and rabbit anti-enolase (Prof Paul Michels, University of Edinburgh) antisera used at 1:1000 or 1:150,000 dilution, respectively, followed by goat antirabbit IRDye™ 800CW antibody (LI-COR) diluted at 1:5,000. Detection of the near infrared signal was monitored using an Odyssey^®^ CLx infrared imaging system (LI-COR). Band signal intensity measurements were carried out using the Analysis Tool in Image Studio™ Lite (Li-COR Biosciences).

### Susceptibility screens

All growth inhibition assays were carried out in a 96-well plate format (ThermoFisher Scientific). Bloodstream form *T. brucei* in the logarithmic phase of growth were seeded at 1 × 10^4^ ml^−1^ (1 × 10^5^ ml^−1^ for *Tbrad51*Δ and *Tbbrca2Δ*) in 200 μl growth medium containing different concentrations of the compound under study: For UV irradiation, parasites received doses up to 9000 J m^−2^ using a Stratalinker^®^ UV crosslinker (Stratagene). After incubation at 37 °C under a 5 % CO2 atmosphere for 3 days, resazurin (Sigma Aldrich) was added to each well at a final concentration of 12.5 μg ml^−1^ (or 2.5 μg per well). The plates were further incubated at 37 °C under a 5 % CO2 for 6 to 8 hours before measuring the fluorescence of each culture using a Gemini Fluorescent Plate reader (Molecular Devices) set at λEX = 530 nm and λEM = 585 nm with a filter cut off at 550 nm. The change in fluorescence resulting from the reduction of resazurin is proportional to the number of live cells. A drug/treatment concentration that inhibits cell growth by 50% (EC_50_) was established using the non-linear regression tool on GraphPad Prism (GraphPad Software Inc.).

## Acknowledgement

We thank Prof Paul Michels (University of Edinburgh) for the *T. brucei* enolase antibody, Dr Richard McCulloch (University of Glasgow) for the anti-T. *brucei* γH2A antisera and the *T. brucei Tbrad51* and *Tbbrca2* null mutant lines, and Heidi Bowring, Charlotte Chaloner, Mariam Kassir and Shiji Zhao (Queen Mary University of London BSc or MSc students) who undertook their research projects in eth Wilkinson lab and contributed to making some of the DNA vectors/cell lines used in this study. We would like to thank Dr Peter Thorpe (Queen Mary University of London) and Dr Martin Taylor (London School of Hygiene & Tropical Medicine) for valuable discussions and comments on the manuscript. Ambika Dittani was funded by Queen Mary University of London College Studentship.

## Supporting information

**S1 Figure: Gene disruption vectors: Construction and their effects on the *T. brucei* genome. (A)**. Sequences corresponding to 5’ and 3’ untranslated (UTR) or coding (CDS) regions of the target gene were amplified using the specified primer combinations (see Table S3) from *T. brucei* genomic DNA. The resultant fragments (sizes in bp) were sequentially cloned either side of a resistance cassette that includes a gene encoding for a *T. brucei* selectable marker (hygromycin B phosphotransferase, neomycin phosphotransferase, blasticidin-S-deaminase or puromycin N-acetyltransferase) plus the *T. brucei* βα or αβ tubulin intergenic repeats (tub IR) required for processing the mRNAs (hatched boxes). The restriction sites used to remove the *T. brucei* integration cassette from the plasmid backbone are noted. (B-G). Schematic representations of the *Tbcsb* **(B)**, Tbexo1 **(C)**, *Tbmre11* **(D)**, Tbsnm1 **(E)**, Tbrad5 **(F)** and Tbrev3 **(G)** alleles and the effects of their disruption with DNA fragments containing sequences encoding for neomycin phosphotransferase (*neo*), hygromycin B phosphotransferase *(hyg)*, blasticidin-S-deaminase *(bla)* or puromycin N-acetyltransferase (*pac*), plus the *T. brucei* tubulin intergenic elements required for processing their mRNAs (hatched boxes). In each panel, P1 and P2 (in black) correspond to the primers used to generate gene specific amplicons from genomic DNA templates while P3, P4, P5 and P6 correspond to *neo, hyg, bla* and *pac* specific primers, respectively: The size of the predicted amplicons, denoted by a dashed line (in kb) is given. P2 and P7 (in grey) correspond to the primers used to generate gene specific amplicons from cDNA templates: The size of the predicted amplicons (in bp) is given. The primer sequences and combinations used for each amplification are listed in Supplementary Tables S2 and S3, respectively.

**S2 Figure. Phenotypic analysis of *T. brucei* null mutants. (A)**. The cumulative cell density of *T. brucei* Tb*exo1*Δ, *Tbmre11* Δ, Tb*csb*Δ, Tbrad5Δ and *Tbrev3*Δ (dashed line) cultures was followed for 8 days and compared against wild type *T. brucei* (solid line) cultures grown in parallel. Each data point represents the mean cell density ± standard deviation from three independent cultures. **(B)**. Cell cycle analysis of *T. brucei* null mutants. The ratio of DAPI-stain kinetoplast (K) and one nuclear (N) genomes in one trypanosomes represents a frequently used marker for the *T. brucei* cell cycle. Phase and DAPI images show cells with a 1K1N, 2K1N and 2K2N arrangement which is indicative of the G1/S, G2/M and post M stages, respectively. The growth shows the relative number of nuclear and mitochondrial genomes structures from asynchronous cultures of wild type, Tb*mre11*Δ Tb*snm1*Δ and Tb*csb*Δ parasites. The number of cells analysed per cell line is given above each bar. **(C)**. The susceptibility of *T. brucei* wild type (solid line) and Tb*mre11*Δ (dotted line) to hydroxyurea, UV, phleomycin and MMS was assessed. All data points are mean values ± standard deviations from experiments performed in quadruplicate. (D). The susceptibility of *T. brucei* wild type and the various null mutant lines against hydroxyurea (HU), phleomycin (Phleo), methyl methanesulfonate (MMS) and ultraviolet radiation (UV), as judged by their EC_50_ values, was compared and expressed as a fold difference.

**S3 Figure. Susceptibility of *T. brucei* null mutants towards DFMO**. Dose response curves and EC_50_ values (in μM) of *T. brucei* wild type (solid line) and Tb*snm1*Δ **(A)**, Tb*exo1*Δ **(B)**, Tb*mre11*Δ **(C)**, Tb*csb*Δ **(D)**, Tbrad5Δ **(E)** and Tbrev3Δ **(F)** (dotted line) to DFMO. All data points are mean values ± standard deviations from experiments performed in quadruplicate.

**S4 Figure. Susceptibility of *T. brucei Tbrad51* and *Tbbrca2* null mutants towards mechlorethamine**. Dose response curves **(A)**. and EC_50_ values (in μM) **(B)** of *T. brucei* wild type (●), Tbrad51Δ (▲) and *Tbbrca2* Δ (○) towards mechlorethamine. All data points are mean values ± standard deviations from experiments performed in quadruplicate. (C). The susceptibility of the *T. brucei* single and double null mutant lines against mechlorethamine, as judged by their EC_50_ values, were compared and expressed as a fold difference relative to wild type.

**S1 Table. *T. brucei* lines used in this project.**

**S2 Table. Primers used in this study.**

**S3 Table. Primers: Function and Combinations.**

